# Modulation of antibiotic sensitivity and biofilm formation in *Pseudomonas aeruginosa* by interspecies diffusible signal factor analogues

**DOI:** 10.1101/291260

**Authors:** Shi-qi An, Julie Murtagh, Kate B. Twomey, Manoj K. Gupta, Timothy P. O’Sullivan, Rebecca Ingram, Miguel A. Valvano, Ji-liang Tang

**Affiliations:** The Wellcome-Wolfson Institute for Experimental Medicine, Queen’s University of Belfast, Belfast BT9 7BL, UK.; School of Microbiology, Biosciences Institute, University College Cork, Cork, Ireland.; School of Chemistry, University College Cork, Cork, Ireland.; School of Pharmacy, University College Cork, Cork, Ireland.; State Key Laboratory for Conservation and Utilization of Subtropical Agro-bioresources, The Key Laboratory of Ministry of Education for Microbial and Plant Genetic Engineering, and College of Life Science and Technology, Guangxi University, Guangxi 530004, China

**Keywords:** Polybacterial infection, cell-cell signaling, chronic infection, antibiotic resistance

## Abstract

The opportunistic pathogen *Pseudomonas aeruginosa* can participate in inter-species communication through signaling by *cis*-2-unsaturated fatty acids of the diffusible signal factor (DSF) family. Sensing these signals involves the histidine kinase PA1396 and leads to altered biofilm formation and increased tolerance to various antibiotics. Here, we show that the membrane-associated sensory input domain of PA1396 has five trans-membrane helices, two of which are required for DSF sensing. DSF binding is associated with enhanced auto-phosphorylation of PA1396 incorporated into liposomes. Further, we examined the ability of synthetic DSF analogues to modulate or inhibit PA1396 activity. Several of these analogues block the ability of DSF to trigger auto-phosphorylation and gene expression, whereas others act as inverse agonists reducing biofilm formation and antibiotic tolerance, both *in vitro* and in murine infection models. These analogues may thus represent lead compounds for novel adjuvants to improve the efficacy of existing antibiotics.

## INTRODUCTION

Antibiotic resistance, coupled with limited development of new antibiotic agents, poses a significant global threat to public health, underscoring the need to find alternative strategies to fight infection^(1, 2, 3)^. These strategies include targeting the signaling pathways that regulate the synthesis of microbial virulence factors and approaches to improve the efficacy of existing antibiotics. Bacterial cell-cell communication (quorum sensing), biofilm formation and cyclic di-GMP signaling were proposed as potential targets for interference by small molecules^(1–4)^. Anti-virulence factors are attractive since they do not influence bacterial growth, which may reduce the selective pressure for developing resistance.

This study addresses the modulation of *Pseudomonas aeruginosa* by inter-species signaling. *P. aeruginosa*, a widespread opportunistic human pathogen, uses two *N*-acyl homoserine lactones (3-oxo-dodecanoyl- and butanoyl-HSL) as well as the quinolone signal PQS (2-heptyl-3-hydroxy-4(1*H*)-quinolone) as quorum sensing molecules that regulate the synthesis of many virulence factors^(5–8)^. There are various reports of small molecule inhibition of these pathways, with consequent effects on virulence factor synthesis^(4, 9–13)^. In addition to intra-species signaling, *P. aeruginosa* can participate in inter-species signaling mediated by molecules of the diffusible signal factor (DSF) family, which are *cis*-2-unsaturated fatty acids. In particular, *P. aeruginosa* senses *cis*-11-methyl-2-dodecenoic acid (DSF) and *cis*-2-dodecenoic acid (BDSF), which are produced by other bacteria such as *Burkholderia* species and *Stenotrophomonas maltophilia* but not by *P. aeruginosa* ^(14, 15)^. Sensing occurs through a histidine kinase PA1396 and leads to altered biofilm formation and increased tolerance to several antibiotics including polymyxin B^(14, 16)^. Mutation of *PA1396* in the model strain PAO1 and several clinical isolates also leads to increased tolerance to polymyxins B and E, suggesting DSF negatively modulates PA1396 activity^(14, 16)^. Detection of DSF family molecules in polybacterial infections, such as those associated with cystic fibrosis (CF) where *P. aeruginosa* is present together with other bacterial species, suggests interspecies signaling occurs *in vivo* and may therefore lead to reduced efficacy of antibiotic therapy^(14, 16)^.

Here, we examine in more detail the role of PA1396 in sensing DSF family signals and the potential of structural analogues of these signals to modulate PA1396 action. In particular, we focus on the effects of DSF analogues on the PA1396-regulated functions of biofilm formation and antibiotic tolerance, both *in vitro* and in murine infection models. We demonstrate that only two of the five trans-membrane helices of the input domain of PA1396 are required for DSF sensing, and DSF binding is associated with enhanced PA1396 auto-phosphorylation. Furthermore, we found that certain analogues of DSF block its ability to trigger auto-phosphorylation, which coincide with specific alterations in the expression of a subset of *P. aeruginosa* genes, reduction in biofilm formation and antibiotic tolerance using both *in vitro* and *in vivo* infection models. Together, our findings indicate DSF-mediated interspecies interactions may be influenced pharmacologically to improve the treatment of chronic *P. aeruginosa* infections.

## RESULTS

### Examining the sensor input domain of PA1396 for DSF recognition

Although DSF perception by PA1396 requires its membrane-associated sensor input domain^(14)^, the underlying molecular mechanism is unknown. As a first step to better understand DSF sensing by PA1396, topological models of the protein were generated with various prediction programs. Analyses with TMHMM, MEMSAT, SOSUI, and HMMTOP suggested a topological model for PA1396 comprising five transmembrane helices (TMHs), and with the N- and the C-terminus located in the periplasm and cytoplasm, respectively (Fig. 1a). In contrast, TOPPRED predicted a relatively large cytosolic loop and a different location in the protein for some of the TMHs. To experimentally assess the PA1396 topology, we constructed translational fusions expressing hybrid proteins of the PA1396 input domain with either alkaline phosphatase (PhoA) or β-galactosidase (LacZ) reporters as topological probes for periplasmic and cytoplasmic locations, respectively. The hybrid proteins were designed such that at least one fusion was placed in each predicted loop facing either the periplasm or the cytoplasm (Fig. 1b).

**Figure 1.**
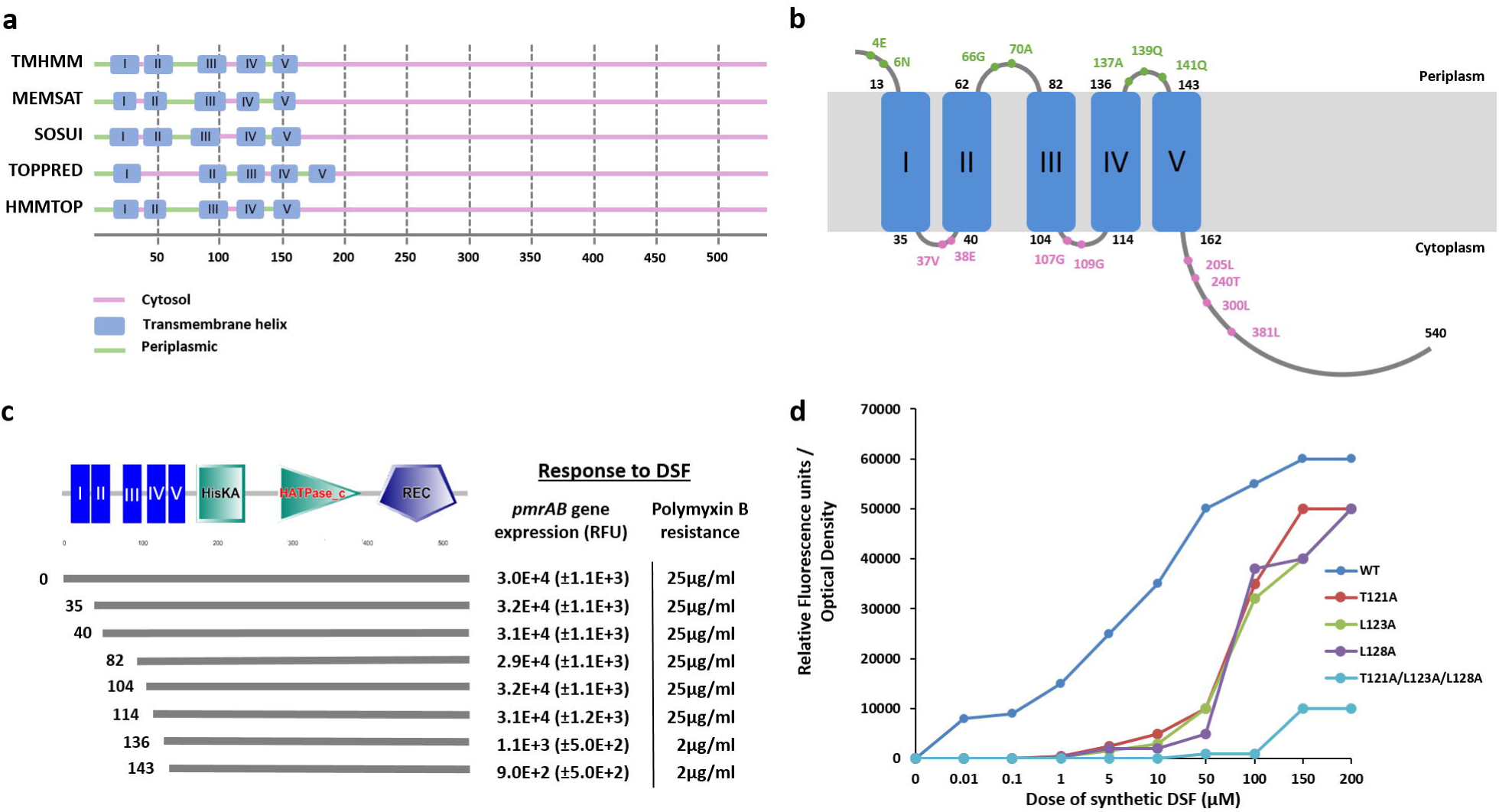
Membrane topology, trans-membrane helices and key residues of sensor kinase PA1396 involved in DSF perception. (a) Graphical representation of the topological predictions of the position of trans-membrane helices (TMHs) of PA1396 made by various membrane topology prediction programs. The location of soluble segments (cytosolic or periplasmic) and the positions of predicted TMHs are indicated. TMHs are designated by Roman numerals. (b) A model of membrane topology of PA1396 derived from reporter fusion data. Phenotypes of *phoA* or *lacZα* reporter fusions together with the amino acid position at which the reporters were fused are indicated by Green (PhoA activity) and Pink (LacZ activity). Numbers next to each predicted TMH indicate the position of amino acid residues at the predicted boundaries between soluble and membrane-embedded regions. (c) Domains of PA1396 as predicted by SMART. Domain abbreviations are: HisKA: His Kinase A (phosphoacceptor) domain; HATPase_c: Histidine kinase-like ATPases; REC: CheY-homologous receiver domain. Blue bars with Roman numerals indicate different transmembrane helices. The lower part of the Figure indicates the effect of progressive N-terminal truncation of PA1396 on the ability of the sensor to respond to DSF, as measured by activation of *pmrAB* gene expression and resistance to polymyxin B. DNA fragments expressing different His6-tagged constructs indicated by the black lines were cloned into pBBR1MCS vector and introduced into the *PA1396* mutant strain. Response to DSF and antibiotic tolerance are reported. (d) Dose–response relationships for activation of a *pmr-gfp* transcriptional fusion by DSF in *P. aeruginosa* strains expressing either wild-type PA1396 or representative variants. The data shown is of a single experiment of three repeated experiments which showed the same trend.

Plasmids carrying hybrid constructs were mobilized into *E. coli* and tested for PhoA and LacZ activity in solid growth medium containing the substrates for either PhoA (X-Phos) or LacZ (X-Gal), which permitted the visual estimation of the enzyme activity of PA1396-PhoA and PA1396-LacZ fusion proteins, respectively. *E. coli* expressing PhoA fusions to PA1396 at amino acid residues E4, N6, G66, A70, A137, Q139 and Q141 exhibited clear PhoA^+^ phenotypes (blue colonies), whereas PhoA fusions to PA1396 at amino acid residues V37, E38, G107, G109 and L205 gave rise to PhoA^−^ phenotypes (white colonies). Conversely, bacteria expressing LacZ fusions to PA1396 at amino acid residues V37, E38, G107, G109, L205, T240, L300 and L381 exhibited LacZ^+^ phenotypes (blue colonies). In contrast, LacZ fusions to PA1396 at amino acid residues E4, N6, G66, A70, A137, Q139 and Q141 resulted in LacZ^−^ phenotypes (white colonies). These findings, also confirmed by quantitative assays (Supplementary Table 1), fit with a model of a five-TMHs protein and the topology shown in Fig. 1b.

To test whether the predicted TMHs are important for DSF sensing, we constructed PA1396 derivatives with progressive truncations at the N-terminus. Each truncated *PA1396* gene was cloned into pBBR1MCS and the constructs introduced into a *PA1396* deletion mutant. Bacteria expressing each of these transconjugants were examined for their ability to respond to exogenous DSF by measuring *pmrAB* gene expression using a *pmr-gfp* fusion, and for polymyxin B resistance (see Methods for details). Removal of the first three TMHs did not affect the ability of PA1396 to respond to DSF when compared to wild-type (Fig. 1c). In contrast, any further truncation led to a loss of DSF responsiveness (Fig. 1c). Western blot analysis with antisera against the His_6_ tag showed that all of the truncated proteins were expressed to the same level as the wild type (Supplementary Fig. 1). These results suggested TMHs IV and V have a role in DSF sensing by PA1396.

To identify specific residues important for DSF binding and signal transduction, we first focused on those conserved in PA1396 homologues from different bacterial species. Conserved or semi-conserved residues in TMHs IV and V (Supplementary Fig. 2) were examined for functionality by constructing alanine replacements (or serine in the case of alanine residues). Residues Y116, L117, T121, L123, L128, T135, P136, W138, A140, P142, M144, L148, M149, V154, I155, P156 and Y158 were replaced and the resulting variants assayed for their ability to confer response to DSF. Bacteria expressing many of the variants did not exhibit significant changes in DSF responsiveness. We identified a number of variants exhibiting decreased sensitivity to DSF, such that higher concentrations were required to activate the *pmr-gfp* reporter although significant activation occurred at concentrations in excess of 50 μM. These variants had alanine replacements at T121, L123, L128, P142, L148, M149, V154 and I155. Except for P142, all the variants with reduced DSF sensitivity were altered in positions within the last two TMHs of PA1396. Variants with combinations of alanine substitutions (such as T121A/L123A/ L128A) had an even further reduced sensitivity (Fig. 1d). The observed differences in the responses to DSF by the reporter strain could not be attributed to differential protein expression since all replacements were expressed to the same level as the parental protein (Supplementary Fig. 1). Together, we conclude residues in TMH-IV and TMH-V of PA1396 play a critical role in the recognition of the DSF signal.

**Figure 2.**
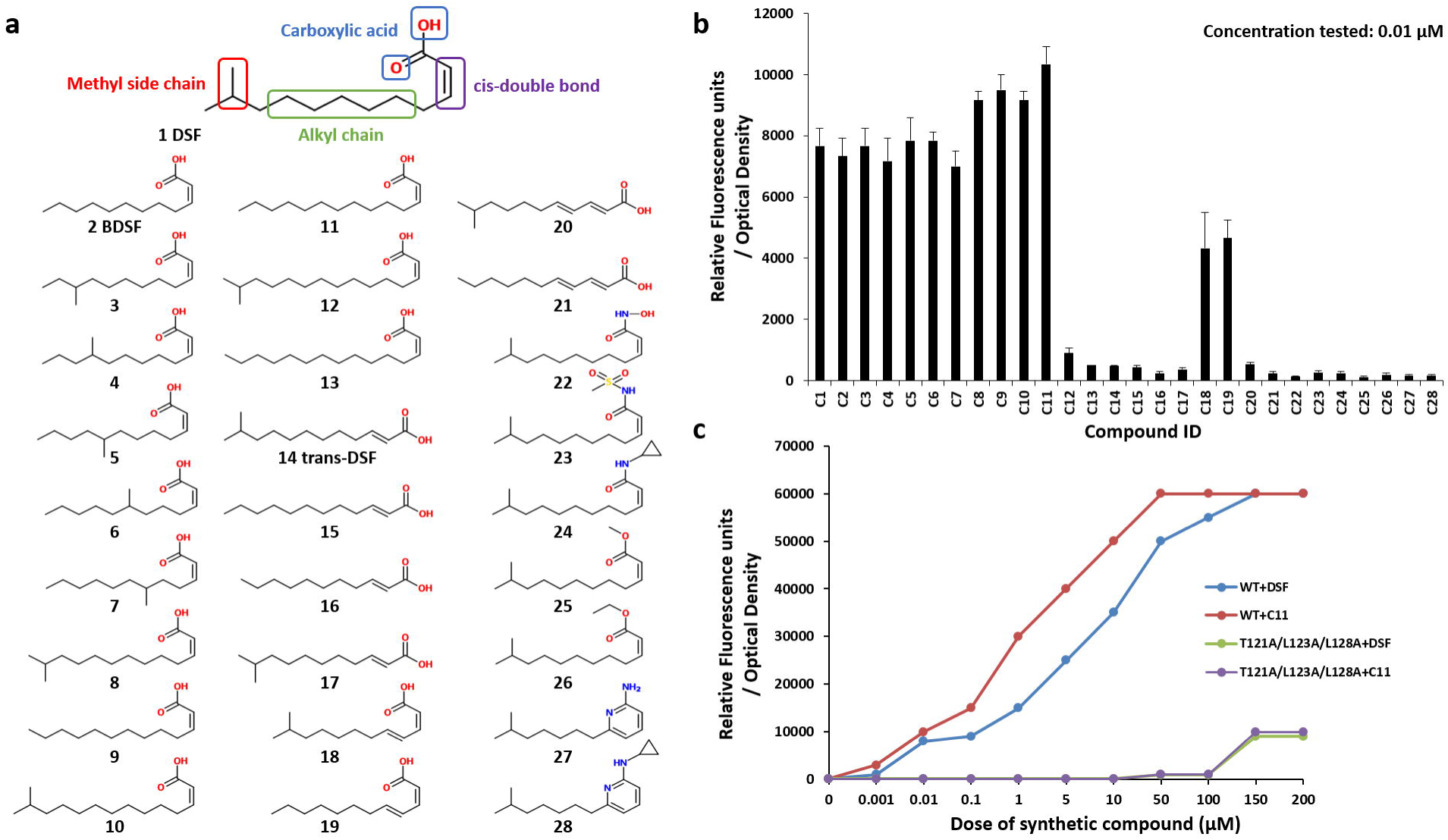
The biological activity of DSF analogues reveals important structural features needed for perception by PA1396. (a) The structure of DSF (*cis*-11-methyl-2-dodecenoic acid) highlighting structural elements. Below this are the chemical structures of the 27 synthetic derivatives of DSF, all validated by ESI-MS and ^1^H NMR spectra as described in the Methods. (b) The biological activity of DSF and its derivatives in activation of the *pmr-gfp* fusion. Samples were tested at a concentration of 0.01 μM (the minimum concentration of *cis*-11-methyl-2-dodecenoic acid required for activation of the bioassay). Values are the average of three repeats (mean± standard deviation). (c) Dose–response of DSF (*cis*-11-methyl-2-dodecenoic acid) and C11 (*cis*-2-tetradecenoic acid) on activation of *pmr-gfp* transcriptional fusion in wild-type PA1396 and a strain expressing the variant T121A/L123A/L128A with alteration in conserved amino acids of TMH IV. The data of GFP fluorescence shown is of a single experiment of three repeats which showed the same trend.

### Physiochemical features of DSF important for recognition by PA1396

The DSF family comprises *cis*-2-unsaturated fatty acids of differing chain length, methyl substitution and saturation^(15)^. Individual bacteria can produce multiple DSF family signals, although often a single chemical species predominates. Previous work showed that the configurational isomer *trans*-11-methyl-2-dodecenoic acid was 500-fold less effective than DSF in activation of *pmrAB*, whereas the corresponding unsaturated fatty acids have little or no activity^(14)^. To examine the structural features of the DSF signals important for PA1396 sensing in more detail, we synthesized a panel of analogs differing from the DSF ‘parent’ molecule in chain length, position and presence of methyl branching or with a *trans* configuration of the double bond. Additional molecules were created by esterification, conversion to hydroxamic acids or amides, and by incorporation of stable aromatic substituents to replace the carboxylic acid moieties (see Fig. 2a). This library of structural derivatives provided a platform to define key structural features important for PA1396 sensing and signal transduction activity.

The activity of the molecules in this panel was assayed by determining the relative fluorescence of the reporter normalized by bacterial cell density (Fig. 2b). Because the minimum concentration of DSF (*cis*-11-methyl-2-dodecenoic acid) for significant induction of *pmr-gfp* expression was 0.01 μM, the 27 DSF derivatives were examined at this concentration. The results revealed that all fatty acids with a *trans* configuration of the double bond at the 2-position render the molecule inactive (Fig. 2b), consistent with previous reports indicating this configuration is critical for DSF activity ^(16–18)^. However, the presence or the position of the methyl group at C-11 in DSF was not essential for activity. Indeed, *cis-*2-tridecenoic acid (C9) and *cis*-2-dodecenoic acid (C2, BDSF) without the methyl substitution, as well as *cis*-7-methyl-2-dodecenoic acid (C6) and *cis*-8-methyl-2-dodecenoic acid (C5) where the methyl group was transposed to the C-7 and C-8 positions, were equally effective in the bioassay. In contrast, the fatty acid chain length played a major role in determining bioactivity. *cis*-2-Unsaturated fatty acids with chain lengths of 12-14 carbons were active whereas those with longer chains were not (Fig. 2b). The introduction of a second double bond in the chain also led to reduced activity, while derivatives in which the carboxylic acid was esterified, converted to a hydroxamic acid or amide derivative or replaced with a 2-amino pyridine moiety were all inactive. For each of these derivatives, no activity was seen in the concentration range from 0.01 to 1000 μM. Interestingly, *cis*-2-tetradecenoic acid (C11), which is a naturally occurring analogue first detected in *Xylella fastidiosa*^(17)^, was most effective in activating the bioassay (Fig. 2c). Strains carrying the T121A/L123A/ L128A replacement mutants of PA1396 showed similar reduced responsiveness to both C11 and DSF (Fig. 2c).

### DSF analogues interfere with DSF signaling in *P. aeruginosa*

The structural analogues unable to trigger DSF-dependent responses were assessed for inhibitory activity against DSF signaling. For these experiments, the molecules were screened at 10 μM using *P. aeruginosa* grown with 50 μM DSF. Several compounds, especially C12, C23 and C24 effectively repressed *pmr-gfp* expression (Fig. 3a), possibly acting as PA1396 receptor antagonists. None of the compounds affected planktonic bacterial growth (Supplementary Fig. 3). Dose response assays of each of the three compounds revealed that compound 23 ((*Z*)-11-methyl-*N*-(methylsulfonyl)dodec-2-enamide; C23) was the most potent DSF antagonist showing substantial inhibition at 0.1 μM (Fig. 3b).

**Figure 3.**
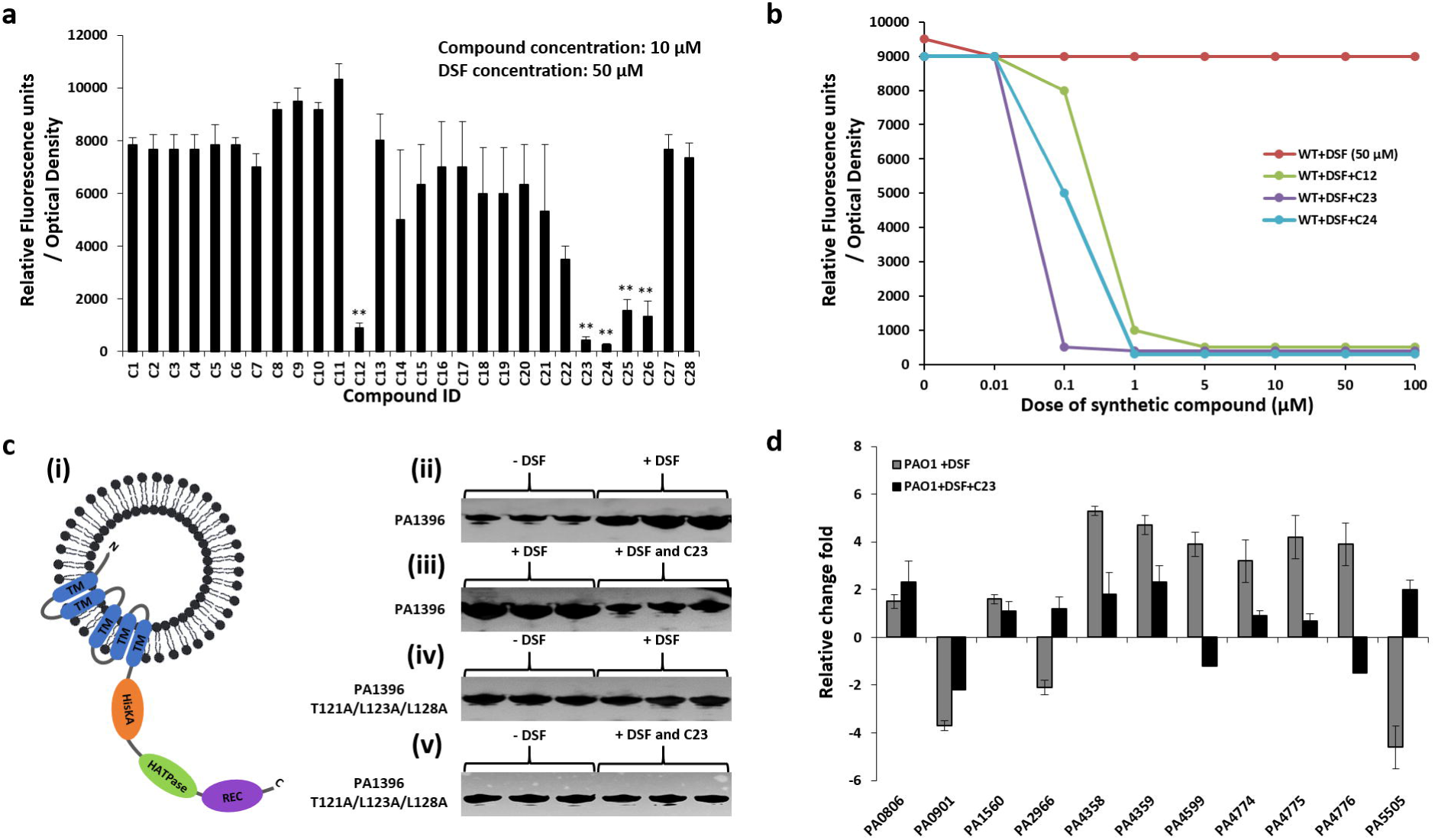
DSF analogues inhibit PA1396 phosphorylation and DSF-dependent activation of genes involved in virulence, antibiotic resistance and biofilm formation. (a) The effects of analogues (at 10 μM) on activation of the *pmr-gfp* transcriptional fusion by DSF (50 μM) was measured as GFP fluorescence. Data shown are means of three replicates and error bars indicate the standard deviations. The asterisks indicate those analogues for which the level of *gfp* expression was significantly different from that of DSF alone (**p < 0.01 as determined by using the Student’ t test). (b) Dose–response relationships of the effects of selected analogues on DSF activation of the *pmr-gfp* transcriptional fusion in wild-type *Pseudomonas aeruginosa*. DSF was present at 50 μM. The analogues tested were C12 (*cis-*13-methyl tetradecenoic acid), C23 (*cis-*11-methyl dodecenoyl methyl sulphonamide) and C24 (*cis-*11-methyl dodecenoyl cyclopropanamide). The data of GFP fluorescence shown is of a single experiment of three repeats, which showed the same trend. (c) PA1396 auto-phosphorylation in response to DSF and analogues. (i) Schematic representation of the orientation of PA1396 in the liposome assay. Domain abbreviations are: HisKA: His Kinase A (phosphoacceptor) domain; HATPase_c: Histidine kinase-like ATPases; REC: CheY-homologous receiver domain. Blue bars indicate different transmembrane helices. (ii) PA1396 auto-phosphorylates in the liposome (lanes 1-3; which are triplicate assays), but increased phosphorylation is seen in response to 10 μM DSF (lanes 4-6); (iii) PA1396 phosphorylation in response to 10 μM DSF (lanes 1-3) is abrogated in the presence of C23 (lanes 4-6); (iv, v) PA1396 (T121A/L123A/L128A) variant protein auto-phosphorylates but does not response to the presence of DSF and/or C23. (d) Differential expression of selected genes implicated in virulence, antibiotic resistance or biofilm formation in the presence DSF, C23 and DSF in combination with C23. Transcript levels of *PA0806*, *PA0901*, *PA1560*, *PA2966*, *PA4358*, *PA4359*, *PA4599*, *PA4774*, *PA4775*, *PA4776* and *PA5505* were examined. The qRT–PCR data were normalised to 16S rRNA and are presented as the fold change with respect to the wild type for each gene. Data (means ± standard deviation) are from three independent biological experiments.

To examine if these compounds interfere directly with signal transduction through PA1396, we assessed the effects of DSF and the putative antagonists on PA1396 auto-phosphorylation. For these experiments, MycHis-tagged PA1396 was expressed, purified under native conditions, and reconstituted in liposomes where the protein adopts an inside-out orientation in which the cytoplasmic histidine acceptor, histidine kinase and receiver domains are surface exposed. The incorporation of PA1396 into liposomes was confirmed by Western blot using anti-Myc antibody, whereas the inside-out orientation can be surmised from the accessibility of ATP to the kinase site, which allows auto-phosphorylation without disruption of the liposomes. Basal auto-phosphorylation of PA1396 in the liposomes was seen in the absence of any signal molecule (Fig. 3c, Supplementary Table 2). However, the addition of DSF to PA1396-loaded liposomes increased the level of auto-phosphorylation dramatically, an effect not seen in the presence of C23 (Fig. 3c, Supplementary Table 2). These findings suggest C23 blocks the recognition of DSF by PA1396, which normally mediates increased kinase activity. The observation that the T121A/L123A/L128A mutant protein had a similar level of basal auto-phosphorylation as the parental PA1396, which could not increase upon addition of DSF, also supports this conclusion (Fig. 3c, Supplementary Table 2). Further, the latter result agrees with the observation that T121A/L123A/L128A has a substantially reduced response to DSF *in vivo* as measured by the *pmr-gfp* reporter bioassay (Fig. 1d).

We previously found that DSF alters the expression levels of *P. aeruginosa* genes implicated in virulence, biofilm formation and stress tolerance ^(14)^. In this study, we assessed the effect of C23 on DSF-regulated functions by qRT-PCR using a subset of 11 DSF-regulated genes. These included genes involved in iron uptake (*PA4358*, *PA4359*) and antibiotic resistance (*PA4599*, *PA4774*–*PA4777*). For these experiments, *P. aeruginosa* was grown in artificial CF sputum medium with and without DSF supplementation, and with both DSF and C23. Cultures were assayed at early log phase (OD_600_ of 0.6). For several of the genes tested, the addition of C23 in the growth medium reduced the level of expression seen with DSF alone and in some cases (e.g. *PA2966, PA5505*) the effects on expression were reversed (Fig. 3d). These experiments were repeated with the *PA1396* mutant strain, demonstrating that the addition of DSF or C23 had no effect on the level of gene expression (Supplementary Fig. 4). Together, the results from *in vitro* auto-phosphorylation experiments and transcriptional profiling of a subset of PA1396-regulated genes indicate C23 interferes with the PA1396 ability to sense DSF.

**Figure 4.**
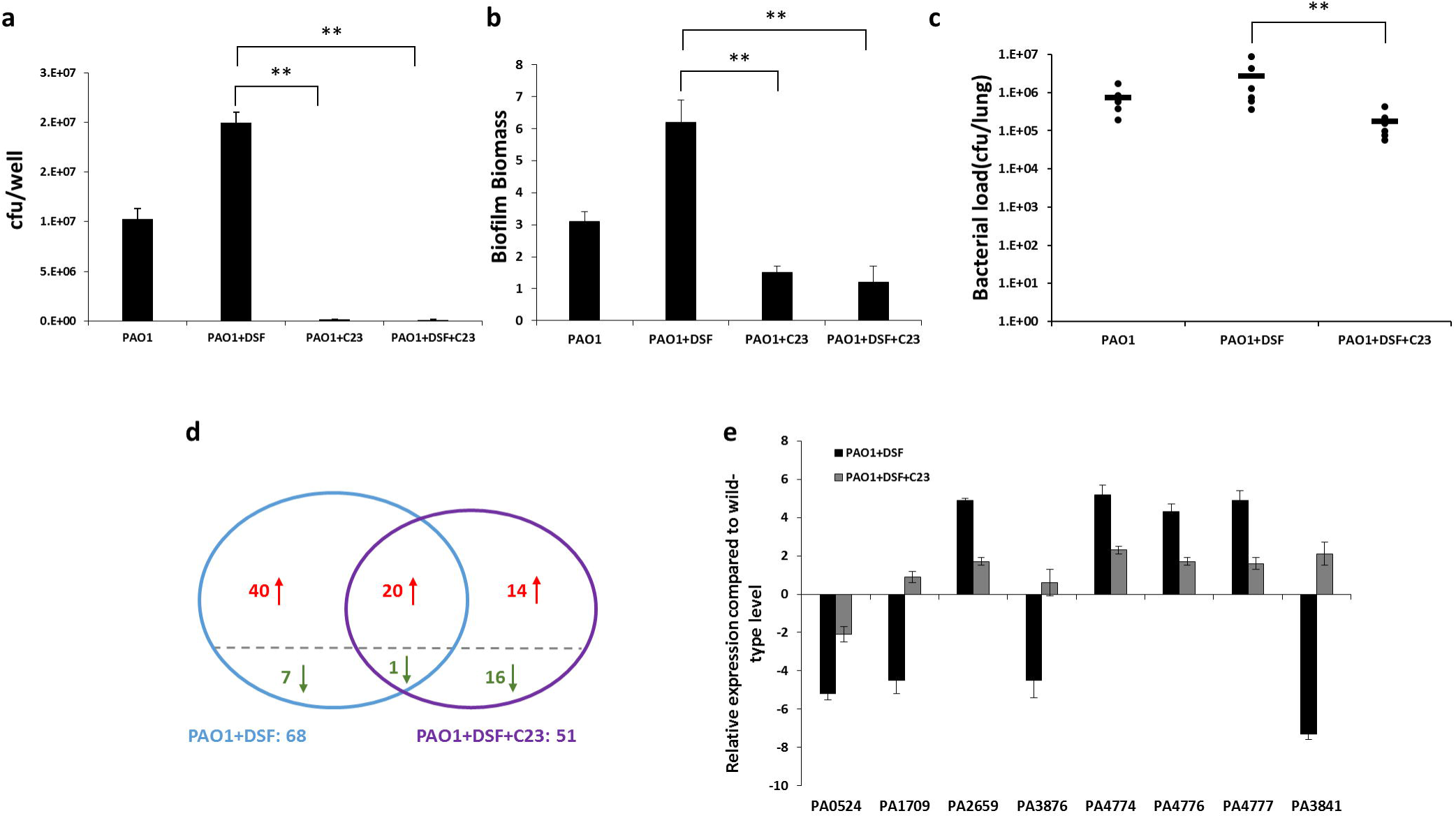
Effects of analogues on DSF-induced biofilm formation and persistence of *P. aeruginosa* during infection. (a) Effect of DSF and C23 alone or in combination on attachment of *P. aeruginosa* strains to CFBE epithelial cells. For these experiments, compounds (0.5 μM) were added to the co-culture at 1h and bacterial attachment to the CFBE epithelial cells was measured after 24 h (see Materials and Methods). (*P<0.01, **p < 0.05, two-tailed Student’s *t*-test). (b) Effect of 0.5 μM DSF and C23 alone or in combination on attachment of *P. aeruginosa* to a glass surface as assessed by crystal violet staining. Biofilm biomass is measured as a ratio of absorbance at 550 and 600 nm. Values given are the mean and standard deviation of triplicate measurements. Asterisks indicate values significantly different from the wild-type (p<0.01, two-tailed Student’s *t*-test). (c) Effect of administration of C23 on *P. aeruginosa* mouse airway infection. C57BL/6 mice were infected intranasally with 1 × 10^7^ CFU of *P. aeruginosa* PAO1 and treated by inhaling PBS with or without 50 μM C23. After 24 h infection, the mice were euthanized, and bacterial loads were determined in lung homogenates. Values represent the mean ± standard deviation. Statistical significance by two tailed Student’s *t*-test is indicated: * *P*<0.05, ** *P*<0.01. (d) Venn diagrams showing results of comparative transcriptome profiling of the effects of addition of either DSF or DSF with C23 on gene expression in wild-type *P. aeruginosa* during mouse lung infection. The comparator is *P. aeruginosa* in mouse lung with no compound addition. Genes significantly up-regulated (> 1.25-fold, P_adj_ < 1 × 10^−5^) are indicated in red, those significantly down-regulated are indicated in green. The complete set of regulated genes is depicted in Supplementary Table 3. (e) qRT-PCR analysis of expression levels of selected genes implicated in virulence and biofilm formation in response to addition of DSF or DSF and C23 in wild-type *P. aeruginosa* during mouse lung infection. Transcript levels of *PA0524*, *PA1709*, *PA2659*, *PA3876*, *PA4774*, *PA4776*, *PA4777* and *PA3841* were examined. The qRT–PCR data were normalised to 16S rRNA and are presented as the fold change with respect to the wild-type for each gene. Data (means ± standard deviation) are representative of three independent biological experiments.

### Interference with DSF sensing decreases biofilm formation and alters virulence of *P. aeruginosa in vivo*

DSF stimulates the development of *Pseudomonas aeruginosa* biofilms *in vitro* and promotes persistent infection *in vivo*^(14, 16, 18)^. We examined if C23 interferes with biofilm formation using a co-culture system in which *P. aeruginosa* biofilms were allowed to form on human lung epithelial cells upon addition of 1 μM DSF^(16)^. C23 at concentrations as low as 0.5 μM C23 led to reduced biofilm formation in the presence of DSF (Fig. 4a). Intriguingly, addition of C23 alone caused a reduction of biofilm development (Fig. 4a). In contrast, the *PA1396* mutant formed more biofilm than the parental strain, but biofilm formation was not significantly altered by either DSF or C23 (Supplementary Fig. 5a). Biofilm formation on a glass surface was also assayed by crystal violet staining. Similar effects of DSF and C23 on biofilm biomass as in the co-culture experiments were observed (Fig. 4b) including the ability of C23 alone to inhibit biofilm formation. These effects were not seen in experiments using the *PA1396* mutant strain (Supplementary Fig. 5b).

**Figure 5.**
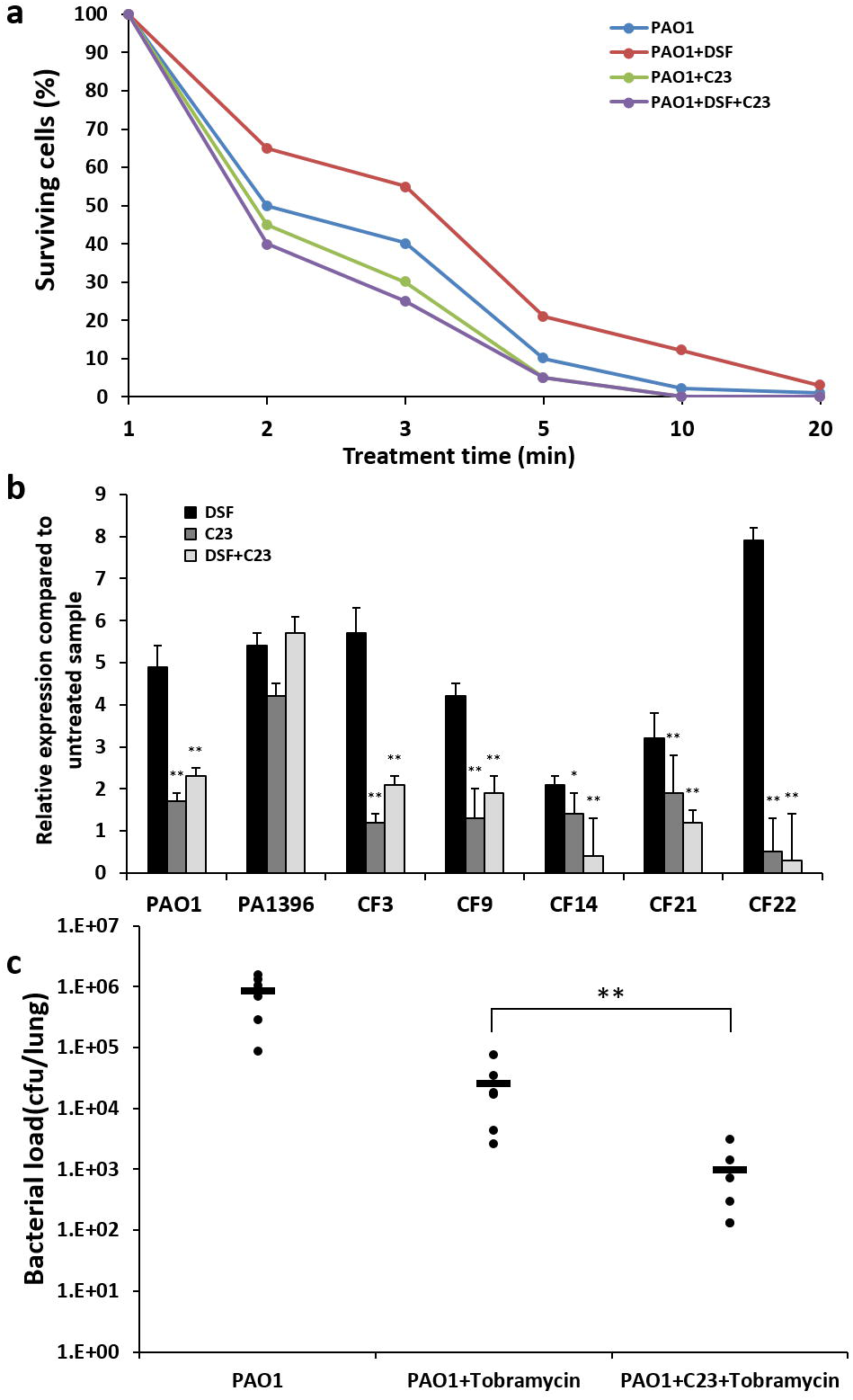
Effects of the DSF analogue C23 on antibiotic resistance of *P. aeruginosa* both *in vitro* and during infection. (a) Effect of addition of DSF, C23 and DSF together with C23 on tolerance of *P. aeruginosa* to polymyxins. Time-courses of killing of *P. aeruginosa* by 4 μg ml^−1^ polymyxin B were established for bacteria suspended in sodium phosphate buffer. Bacteria for these experiments were grown in BM2 medium with glucose supplemented with 2 mM Mg^2+^. DSF and C23 were added to these cultures to a final concentration of 50 μM. The data shown is of a single experiment of three repeated experiments which showed the same trend. (b) Effect of addition of DSF, C23 and DSF with C23 on antibiotic resistance gene expression in clinical isolates of *P. aeruginosa* as compared to the laboratory strain PAO1. Transcript levels of *PA4775* in PAO1, the isogenic *PA1396* mutant and a selection of clinical isolates of *P. aeruginosa* (CF3, CF9, CF14, CF21 and CF 22) were examined using qRT-PCR as described in Methods. The qRT–PCR data were normalised to 16S rRNA and are presented as the fold change relative to the wild-type for each gene in each strain with no addition. Data (means ± standard deviation) are representative of three independent biological experiments. The statistical significance of the difference between DSF alone and C23 plus DSF or C23 alone was examined by two tailed Student’s *t*-test and are indicated as: *: *P*<0.05; **: *P*<0.01. (c) Effect of administration of C23 on *P. aeruginosa* clearance by tobramycin treatment in the mouse airway. C57BL/6 mice were infected intranasally with 1 × 10^7^ CFU *P. aeruginosa* PAO1 and treated by inhaling PBS with or without 50 μM C23. Tobramycin was administered at a concentration of 30 mg/kg 1 h after infection. After 24 h of infection, the mice were harvested, and bacterial loads were determined in lung homogenates. Values represent the mean ± standard deviation. Statistical significance of the difference between C23 plus tobramycin treatment and tobramycin alone was examined by two tailed Student’s *t*-test and are indicated: **: *P*<0.01.

To determine whether C23 modulates bacterial behavior during infections, C57BL/6 mice were inoculated intranasally with suspensions of *P. aeruginosa* supplemented with C23, DSF or C23 with DSF. As a control, bacterial inocula were suspended in PBS alone. At 24 h post infection, the *P. aeruginosa* PAO1 bacterial load was quite considerable in the control animals. However, in the presence of C23, the number of bacteria was considerably reduced. With DSF alone, higher numbers of bacteria were seen after 24 h than in the control (Fig. 4c), in agreement with previous trends seen in other *in vivo* models ^(16)^. With C23 in combination with DSF, the bacterial load was reduced when compared to DSF alone (Fig. 4c). Thus, C23 reduced the bacterial load in this mouse model in the presence or absence of DSF.

To verify that C23 functions by inhibiting DSF signaling *in vivo*, we used microarray to compare the effects of adding either DSF alone or DSF with C23 on *P. aeruginosa* gene expression during mouse lung infection relative to the gene expression of *P. aeruginosa* in the mouse lung with no compound. RNA was derived from organisms isolated directly from the lung homogenates of infected mice. The results showed that the expression of 68 genes was significantly altered (> 1.25-fold, P_adj_ < 1 × 10^−5^) by the addition of DSF (Fig. 4d; Supplementary Table 3). While many of these differentially expressed genes were originally annotated as hypothetical proteins, several encode factors required by *P. aeruginosa* for growth and/or host colonization of the host; these genes included *norB* (*PA0524*), *popD* (*PA1709*), *sphA* (PA2659) and *narK* (*PA3876*). Additional regulated genes encoded factors that contribute to *P. aeruginosa* virulence and biofilm formation such as the *mexC* (*PA4599*), *pmr* operon (*PA4774, PA4776, PA4777*) and *exoS* (*PA3841*).

Addition of DSF and C23 significantly altered expression of 51 genes (Fig. 4d). Comparison with the effects of DSF alone showed expression changes in 21 genes occurred in both treatments (DSF vs DSF plus C23) with all 21 of these genes altered in the same direction. The expression level (fold change in expression) of these genes was reduced in DSF plus C23 treatment when compared with the DSF treatment alone (Supplementary Table 3). Further, some genes significantly regulated by DSF showed no significant alteration in the presence of DSF and C23. Examples of this group of genes include *popD* (*PA1709*) and *pmr* operon genes (*PA4774, PA4776, PA4777*). Also, various genes showed a significant alteration in expression in response to DSF plus C23, but not to DSF alone (Fig. 4d, 4e). Selected genes were examined using qRT-PCR to confirm the alteration in expression as revealed by the microarray (Fig. 4e). Together, the combined results of the experiments described above show that C23 is not only able to inhibit DSF-activated responses *in vitro* and *in vivo*, but also has a broader effect on gene expression independent of DSF-regulated responses.

The effect of C23 alone on gene expression was examined in a separate experiment on bacteria grown in artificial sputum medium in the presence or absence of the compound at 50 μM. RNA was extracted from these cultures and the level of expression of selected genes examined by qRT-PCR. Of the genes examined, *gcdH* (*PA0447*) and *murA* (*PA4450*) appeared to have elevated expression as a response of C23 addition to *P. aeruginosa* PAO1, but showed little or no response to DSF (Supplementary Fig. 6).

### The DSF analogue C23 improves antibiotic efficacy

Perception of DSF by *P. aeruginosa* mediated by PA1396 leads to increased expression of the *pmrAB* operon and the concomitant increased level of resistance to the cationic antimicrobial peptides polymyxin B and E ^(14, 16)^. Further, DSF increases tolerance to polymyxin B in *P. aeruginosa* biofilms-airway epithelial cells co-cultures^(16)^. These observations prompted us to examine the influence of C23 on the sensitivity of *P. aeruginosa* to antibiotics. As expected, addition of DSF led to *pmrAB* up-regulation and an increased resistance to polymyxin B (Fig. 5a, b). By contrast, addition of C23 plus DSF led to a marked reduction in *pmrAB* expression and reduced resistance to polymyxin B. Although the *pmrAB* expression level in response to C23 alone was close to wild-type, C23 did appear to slightly enhance the activity of polymyxin B against PAO1 (Fig. 5a). We also examined the influence of DSF and C23 singly and in combination in resistance to other classes of antibiotics used in treatment of *P. aeruginosa*. Addition of DSF to the cultures led to increased resistance to tobramycin and nalidixic acid but had no effect on carbapenem resistance. The effect of DSF on increased tobramycin and nalidixic acid resistance was reversed by C23 (Supplementary Fig. 7).

To validate these results in other strains, we examined a panel of *P. aeruginosa* clinical isolates and the data revealed that addition of DSF led to a significant increase in the expression of *pmrAB* in each of these strains (Fig. 5b). However, addition of C23 reversed this DSF-induced effect in all strains, whereas C23 alone gave only modest or no increase in expression. In all of these cases, changes in *pmrAB* expression were associated with a concomitant alteration in resistance to polymyxin B, as seen with PAO1 (Supplementary Fig. 7). Further, addition of C23 also reversed DSF-induced tobramycin tolerance in the clinical isolates, which decreased in all strains (Supplementary Fig. 8). The *PA1396* mutant in PAO1 showed increased resistance to polymyxin B, an effect that could not be reversed by addition of C23, as expected. To determine whether PA1396 was also required for the response of clinical isolates of *P. aeruginosa* to C23, we created *PA1396* gene disruption mutants in five of them, which were selected because of their intrinsically higher level of *pmrAB* expression and polymyxin resistance. None of the mutant strains responded to C23, confirming the C23 antagonist effect on DSF is mediated by PA1396 (Supplementary Fig. 9).

We also assessed the effect of C23 on the efficacy of tobramycin treatment in the C57BL/6 mouse lung infection model. Mice were infected with 1 × 10^7^ CFU of PAO1 and treated by inhaling PBS with or without C23. Tobramycin was administered at a concentration of 30 mg/kg one hour after infection, as previously reported^(19, 20)^. The lung bacterial load was determined 24 h post-infection. Control mice infected with *P. aeruginosa* and inhaled PBS exhibited considerable colonization (Fig. 5c), which was reduced by tobramycin treatment. However, adding C23 with tobramycin resulted in a larger decrease of the bacterial load than that seen with tobramycin alone (Fig. 5c). The kinetics of clearance of the *PA1396* mutant strain from mice not treated with antibiotics was similar to that of mice infected with PAO1. Together, these results demonstrate that C23 treatment leads to reduced resistance of *P. aeruginosa* laboratory and clinical strains to distinct antibiotics both *in vitro* and *in vivo* during infection, an effect manifested through a functional PA1396 sensor kinase.

## DISCUSSION

This study provides a detailed characterization of the membrane topology of PA1396 including the identification of its input domain, which has considerable sequence similarity with the input domain of RpfC, the sensor histidine kinase for DSF-mediated cell-to-cell signalling in *Xanthomonas campestris* pv. *campestris* ^(15, 23)^. However, significant differences between PA1396 and RpfC also exist. RpfC contains a short N-terminal periplasmic region implicated in DSF binding^(24)^, suggesting DSF is captured in the periplasmic space. Residue E19, which is conserved in all homologues with the RpfC-related input domain, is among various key residues for DSF binding in RpfC^(14, 24)^. In *P. aeruginosa* PA1396, the corresponding residue is E8, also in the N-terminal periplasmic region. However, deletion of a significant part of the N-terminal region of PA1396 has no effect on DSF sensing by *P. aeruginosa*. Unlike RpfC, our results indicate the DSF binding capability of PA1396 requires residues in TMHs IV and V. This suggests that DSF likely interacts with residues in these TMHs at the membrane level.

The PA1396 recognition of DSF shares features with the CqsS system of *Vibrio cholerae*, which is responsible for detection of the quorum-sensing signal CAI-1 (*S*-3-hydroxytridecanone). Both CqsS and PA1396 have complex membrane spanning domains with multiple TMHs and periplasmic and cytoplasmic loops of limited size and both recognise ligands with alkyl chains of medium length and an amphipathic nature. Nevertheless, substantial differences between the systems occur. In CqsS, conserved residues in the first three (of six) TMHs are obligatory for CAI-1 ligand binding and subsequent signal transduction. Further, residues in the fourth transmembrane helix enable CqsS to discriminate between CAI-1 and amino-CAI-1 and residues in the fifth TMH has roles in restricting ligand head group size and tail length^(25)^. In contrast, PA1396 truncation experiments suggest that the first three TMHs of the protein are dispensable for DSF perception. However, whether THMs I-III of PA1396 have roles in detection of other ligands is unknown.

Previous work on interspecies signaling in *P. aeruginosa* demonstrated that the *cis*-2-unsaturated fatty acids DSF and BDSF trigger *pmr* gene expression, but weak or no activity was observed with the *trans*-derivatives and related saturated fatty acids. Here we confirm and extend the analysis of structural requirements for interspecies signaling. All *trans*-unsaturated fatty derivatives, irrespective of chain length, had little or no signaling activity on *pmr* gene expression. Similarly, any derivative lacking a free carboxylic acid group (through esterification, conversion to a hydroxamic acid or substitution with a 2-amino pyridine moiety) was inactive. The signaling activity was not significantly affected by the position of the methyl group (at position 11 in DSF) but was partially reduced by introducing a second double bond. Notably, a derivative of DSF with a second double bond (*cis, cis*-11-methyldodeca-2,5-dienoic acid) occurs naturally in various *Xanthomonas* species, but also has a reduced ability compared to DSF to trigger DSF-regulated genes^(18)^. Further, interspecies signaling in *P. aeruginosa* depends on the chain length of the *cis*-unsaturated fatty acid; molecules with chain lengths longer than 14 (this work) and less than 12^(15, 18, 21, 22)^ have no activity. In particular, PA1396 does not respond to *cis*-2-decenoic acid, a molecule produced by *P. aeruginosa* and which is implicated in biofilm dispersal ^(15, 23)^.

We demonstrate that DSF and *cis*-unsaturated fatty acid analogues enhance the tolerance of *P. aeruginosa* to a range of antibiotics including polymyxins, tobramycin and nalidixic acid; this action requires the sensor kinase PA1396. In contrast, Deng and colleagues^(26)^ reported that both DSF and its analogues promote sensitivity of several Gram-positive pathogens including *Bacillus cereus* and *Staphylococcus aureus* to various antibiotics including kanamycin and gentamicin. The molecular mechanisms underlying these effects in *B. cereus* are unclear, but it is intriguing that *cis*- and *trans*-2-unsaturated fatty acids are equally effective in inducing the response.

We reasoned that if DSF signaling activates expression of genes involved in antibiotic tolerance, structural analogues of the molecule could act as antagonists. Accordingly, we show that various effects of DSF on *P. aeruginosa* including induction of *pmr* gene expression and persistence in the mouse model is antagonised by the C23 analogue. Antagonism of DSF action may have an application to control of *P. aeruginosa* in multispecies infections with pathogens that produce DSF family signals. An example is infection associated with cystic fibrosis (CF), where *P. aeruginosa* can occur together with *S. maltophilia* and *Burkholderia* species, both of which produce DSF and BDSF. The detection of DSF and BDSF signals at physiologically relevant levels in the sputum of CF patients supports the contention that interspecies signaling may occur in these infections, where it may affect the antibiotic sensitivity of *P. aeruginosa* as suggested by *in vitro* experiments.

Intriguingly, C23 has additional modulating effects on *P. aeruginosa* that cannot be attributed to its role as a DSF antagonist. For example, C23 affects *gcdH* and *murA* expression, two genes that do not respond to DSF. C23 also affects biofilm formation, both in co-culture with lung epithelial cells and in ‘complex’ medium, in DSF-independent manner and contributes to a more rapid clearance of *P. aeruginosa* from the lungs of mice. More detailed molecular knowledge of the activities of PA1396 may afford insight into these different actions of C23. For example, can PA1396 sense other environmental signals for biofilm formation (in addition to DSF) perhaps through TMHs I-III? Is transduction of these signals also blocked by C23? Can C23 alone cause reduced auto-phosphorylation of PA1396 *in vivo*? Our unpublished work indicate the two-component regulator PA1397 is involved in signal transduction beyond PA1396 (S. An, unpublished).However, the findings are not consistent with a simple model in which DSF-induced phosphorylation of PA1397 directly activates *pmr* gene expression, indicating that other regulatory elements are involved. Future work will be directed at identifying these components.

In conclusion, pharmacological inhibition of DSF-mediated interspecies signaling in *P. aeruginosa* using a DSF analogue potential has identified a potential lead compounds for molecules that could be used as antibiotic adjuvants to control diseases caused by this human pathogen.

## MATERIALS AND METHODS

### Bacterial strains and culture conditions

*Pseudomonas aeruginosa* PAO1 was obtained from the Genetic Stock Center (http://www.pseudomonas.med.ecu.edu/)(strain PAO0001). Other bacterial strains and plasmids used in this study are listed in Supplementary Table 4. *P. aeruginosa* and other strains were routinely grown at 37°C in Luria–Bertani (LB) medium while *Xanthomonas campestris* strains were routinely grown at 30°C in NYGB medium, which comprises Bacteriological Peptone (Oxoid, Basingstoke, UK), 5 g l^−1^; yeast extract (Difco), 3 g l^−1^ and glycerol, 20 g l^−1^. The FABL medium consists of 97% FAB medium [(NH_4_)_2_SO_4_, 2 g l^−1^; Na_2_HPO_4_ 2H_2_O, 6 g l^−1^; KH_2_PO_4_, 3 g l^−1^; NaCl, 3 g l^−1^; MgCl_2_, 93 mg l^−1^; CaCl_2_, 11 mg l^−1^) and 3% L medium (Bactotryptone, 10 g l^−1^; yeast extract, 5 g l^−1^; sodium chloride, 5 g l^−1^; and D-glucose 1 g l^−1^). Cultures were also grown in artificial sputum medium which comprises: 5 g mucin from pig stomach mucosa (Sigma), 4 g DNA (Fluka), 5.9 mg diethylene triamine pentaacetic acid (Sigma), 5 g NaCl, 2.2 g KCl, 5 ml egg yolk emulsion (Oxoid) and 5 g amino acids per 1 l water (pH 7.0) ^(16, 19)^. The antibiotics used included tobramycin, polymyxin, kanamycin, rifampicin, gentamycin, spectinomycin, nalidixic acid, carbapenem and tetracycline at the indicated concentrations.

### DNA manipulation

Molecular biological methods such as isolation of plasmid and chromosomal DNA, PCR, plasmid transformation, as well as restriction digestion were carried out using standard protocols. PCR products were cleaned using the Qiaquick PCR purification kit (Qiagen) and DNA fragments were recovered from agarose gels using Qiaquick minielute gel purification kit (Qiagen). Oligonucleotide primers were purchased from Sigma-Genosys. Primer sequences are provided in Supplementary Table 5.

### Cloning the *PA1396* gene

The DNA fragments encoding the full-length PA1396 protein or truncation of interest were synthesized by Gene Oracle (Santa Clara, USA) in pGOv4 and sub-cloned into pET47b, pME6032 or pBAD/Myc-His before transformation into *E. coli* BL21 (DE3). Genomic regions are described in Supplementary Table 5. BL21 (DE3) cells were grown in LB media and induced with 0.25 mM IPTG; protein overexpression was carried out at 37°C for 1 h. Purification was achieved by Ni^2+^ affinity chromatography using the N-terminal His6 tag.

### Construction of targeted *PA1396*–*phoA* and *PA1396*–*lacZ* fusions

Trans membrane domain (TMD) predictions of the *P. aeruginosa* PA1396 protein were obtained using the TOPRED^(27)^, TMHMM^(28)^, DAS-TMFILTER^(29, 30)^ SOSUI^(31)^ and HMMTOP^(32)^ programs, each with their default settings.

Suitable fusion sites in the periplasmic and cytoplasmic loops were identified using a consensus based on the TMS prediction data obtained (Supplementary Fig. 1). The sites selected correspond to amino acids E4, N6, V37, E38, G66, A70, G107, G109, A137, Q139, Q141 and L205. The DNA fragments encoding the proteins of interest were synthesized by Gene Oracle (Santa Clara, USA) in pGOv4 and sub-cloned into topology reporter plasmids *phoA* (pRMCD28) and *lacZ* (pRMCD70). Genomic regions are described in Supplementary Table 5.

The fusion junction of each construct was confirmed by sequencing. Alkaline phosphatase assays were carried out according to Daniels *et al*. (1998) ^(33)^. β-Gal assays were carried out according to Baker *et al.* (1997) ^(34)^, except that overnight cultures were sub-cultured and grown to OD<0.6 before activity assays. Samples were assayed in triplicate over at least three independent experiments.

### Truncation and mutagenesis of PA1396 gene

To construct strains harboring truncated *PA1396* alleles, we employed plasmid pME6032. We amplified *PA1396* alleles with deleted for amino acids 1–35, 1–40, 1– 82, 1–104, 1–114, 1–136 and 1–143 respectively. These amplified fragments were ligated into pME6032 using appropriate restriction sites. Primer sequences and restriction sites are provided in Supplementary Table 5. The DNA fragments encoding the proteins with alterations in Y116, L117, T121, L123, L128, T135, P136, W138, A140, Q142, M144, L148, M149, V154, I155 and F157 were synthesized by Gene Oracle (Santa Clara, USA) in pGOv4 and sub-cloned pME6032. All constructed plasmids were transformed into strain PA1396 mutant selecting for Gm^R^.

### Synthesis of DSF analogues

The starting point for the synthesis of *cis*-2-dodecenoic acid C2 (BDSF) and 11-methyldodec-2-enoic acid C1 (DSF) involved a Swern oxidation of the starting alcohol with dimethyl sulfoxide (DMSO) and oxalyl chloride at −78 °C to corresponding aldehydes decanal and 11-methyldecanal^(35)^. A subsequent Wittig reaction of the purified aldehyde with the modified Horner-Wadsworth-Emmons phosphonate salt ethyl [bis(2,2,2-trifluoroethoxy)-phosphinyl]acetate in the presence of sodium hydride (NaH) in THF afforded both the *cis-* and *trans-*α,β-unsaturated esters, ethyl dodec-2-enoate and ethyl 11-methyldodec-2-enoate^(36)^. Hydrolysis of the *cis*-α,β-unsaturated esters with lithium hydroxide (LiOH) in THF:MeOH:H_2_O (2:1:1 (v/v/v)) gave required products C2 and C1^(37,38)^. Treatment of C1 with propylphosphonic anhydride and cyclopropylamine gave the required amide analogue^(39)^. DSF antagonist C23 was prepared by way of an EDCI-mediated coupling of C1 with methanesulfonamide in the presence of dimethylaminopyridine and subsequent isolation of the desired *cis*-isomer by reverse phase HPLC. All the chemicals and reagents were purchased from Sigma-Aldrich unless otherwise stated. Additional synthetic procedures and analytical data used in this study are delineated in Supplementary Note.

### DSF analogue structure analysis

Nuclear magnetic resonance spectra ^1^H, ^13^C, ^1^H-^1^H COSY and DEPT were recorded on a Bruker Avance 400 NMR Spectrometer (400 MHz for ^1^H) and a Bruker Avance 500 NMR Spectrometer (500 MHz for 1H and 125 MHz for ^13^C) with trimethylsilylchloride as an internal standard in CDCl_3_. ^1^H-^13^C correlated HMBC and HMQC spectra were performed by Bruker Avance 500 spectrometer. Mass spectra ESI-MS and High-Resolution Mass Spectra ESI-MS were performed on a Waters/Micromass: LCT Premier Time of Flight and a Quattro Micro triple quadrupole instruments respectively. Infrared spectra were measured using NaCl plates on a Perkin Elmer paragon 1000 FT-IR spectrometer.

### DSF bioassays

The original DSF bioassay is based on its ability to restore endoglucanase production to *rpfF* mutant of *Xcc* as described in ^(40)^. DSF activity was expressed as the fold increases in endoglucanase activity over the control. We also used the bioassay previously described^(16)^ that relies on DSF-dependent induction of fluorescence (*pmr-gfp*) in *P. aeruginosa* PAO1.

### Reconstitution and phosphorylation of PA1396-His in liposomes

Using a variation in the method previously described^(41,42)^, *E. coli* strains containing pBAD/*Myc*-His (PA1396-pBAD/*Myc*-His) were induced with 0.2% arabinose and purified through nickel columns according to the manufacturer’s instructions (Qiagen). Liposomes were reconstituted as previously described^(24,41)^. Briefly, 50 mg of *E. coli* phospholipids (44 μL of 25 mg/mL; Avanti Polar Lipids) were evaporated and then dissolved into 5 ml of potassium phosphate buffer containing 80 mg of *N-*octyl-β-D-glucopyranoside. The solution was dialyzed overnight against potassium phosphate buffer. The resulting liposome suspension was subjected to freeze–thaw in liquid nitrogen. Liposome size was analyzed by dynamic light scattering. Liposomes were stored at 4 °C. Liposomes were then destabilized by the addition of 26 mg of dodecylmaltoside, and 5 mg of PA1396-His (dual Myc- and His-tagged protein) was added, followed by stirring at room temperature for 10 min.

Two hundred-sixty milligrams of Biobeads (BioRad) were then added to remove the detergent, and the resulting solution was allowed to incubate at 4°C overnight. The supernatant was then incubated with fresh Biobeads (BioRad) for 1 h in the morning. The resulting liposomes containing reconstituted PA1396-His were frozen in liquid N2 and stored at −80°C until used. The orientation of HKs in the liposome system was established by other groups and can be confirmed from the accessibility of ATP to the kinase site and anti-Myc antisera to the C-terminal PA1396-MycTag without disruption of the liposomes.

Twenty microliters of the liposomes containing PA1396-His were adjusted to 10 mM MgCl2 and 1 mM DTT, and various concentrations of agonist or antagonist, frozen and thawed rapidly in liquid nitrogen, and kept at room temperature for 1 h (this allows for the signals to be loaded within the liposomes). [γ32P]dATP (0.625 μl) (110 TBq/mmol) was added to each reaction. To some reactions, 10 μg of PA1396-His was added. At each time point (0, 10, 30, 60, or 120 min), 20 μl of SDS loading buffer was added. For all experiments involving PA1396 alone, a time point of 10 min was used. The samples were run on SDS/PAGE without boiling and visualized using a Molecular Dynamics PhosphorImager. These were quantified by ImageJ software.

### Biofilm Assays

Biofilm was assessed by attachment to glass and was determined by crystal violet staining. Log-phase-grown bacteria were diluted to OD_600_ nm = 0.02 in LB broth and 5 ml was incubated at 37°C for 24 h in 14-ml glass tubes. After gently pouring off the media, bacterial pellicles were wash ed twice with water and were then stained with 0.1% crystal violet. Tubes were washed and rinsed with water until all unbound dye was removed. Bound crystal violet was eluted in ethanol and measured at OD_595_. Three independent assays were carried out for each strain.

### Polymyxin killing curves

Killing curves were carried out at 37°C temperature as previously described^(14, 16)^. In short, mid-log phase cultures (OD_600_ 0.4–0.6) of PAO1 wild-type strain, PAO1 supplemented with 50 μM DSF or PA1396 mutant grown in BM2 medium supplemented with 0.5 mM and 2 mM Mg^2+^ were diluted 1:100 into 30 mM sodium phosphate buffer, pH 7.0, containing 4 μg ml^−1^ of polymyxin B sulphate and 5 μg ml^−1^ polymyxin E sulphate (Sigma). Samples were shaken gently, and aliquots removed at specified time intervals were assayed for survivors by plating appropriate dilutions onto LB agar.

### Infection and treatment of animals

All animal experiments were approved by the Animal Ethics Committees of Queens College Belfast and Nanyang Technological University. Mouse infection was performed using a variation on the mouse model described in ^(16, 43)^. Briefly, *P. aeruginosa* strains were grown in Luria broth at 37 °C overnight with shaking, after which bacteria were collected by centrifugation and resuspended in PBS. The exact number of bacteria was determined by plating serial dilutions of each inoculum on pseudomonas isolation agar plates. Female C57BL/6 mice (approximately 8-weeks old) were anesthetized and intranasally inoculated with 20 μl of the bacterial suspension (1 × 10^7^ CFU/mouse) in PBS with PAO1, PAO1/DSF, PAO1/or selected DSF analogue or *PA1396* mutant. Mice were killed at 24 h post-infection by intraperitoneal injection of 0.3 ml of 30% pentobarbital. Lungs were harvested aseptically and homogenized in sterile PBS. A 10-fold serial dilution of lung homogenates was plated on *Pseudomonas* isolation agar. The results (means±s.d.) are expressed as CFU/lung.

For animal experiments examining the effectiveness of tobramycin treatment against *P. aeruginosa* during respiratory infection, the mice were intranasally inoculated with 20 μl of the bacterial suspension in PBS with PAO1, PAO1/DSF, PAO1/or selected DSF analogue (1 × 10^7^ CFU/mouse). Each mouse received was administered tobramycin 1 h post-infection. 24 h after infection, the lungs were removed, homogenized with PBS and plated on agar to determine the number of viable bacteria. The results (means±s.d.) are expressed as CFU/lung.

### Gene expression-profiling experiments

For the *in vivo* RNA samples, infected animals as described above were euthanized ~24 h post-infection. RNA was isolated from airway homogenate pellets using Trizol according to the manufacturer’s instructions. Samples were treated with DNAse, then rRNA was removed from 1 to 5 μg of total RNA using the RiboZero kit for Gram-negative bacteria (Epicentre). rRNA-depleted samples were concentrated by EtOH precipitation. cDNA synthesis and hybridization to Affymetrix GeneChip *P. aeruginosa* genome arrays were carried out by a commercial Affymetrix Genechip service supplier (Conway Institute of Biomolecular & Biomedical Research, UCD, Ireland). Microarray data analysis was performed using Biomedical Genomics Workbench software (Qiagen).

### Quantitative real-time PCR assays

*P. aeruginosa* strains (including mutants) were inoculated at an OD_600_ of 0.1 in LB (with 50 μM DSF-like signal molecules if used). Cultures were harvested during mid-log phase (OD_600_ = 0.6) and RNA was extracted using the High Pure RNA Isolation Kit (Roche). The expression of genes was monitored by qRT-PCR, as described previously ^(44,45)^.

### Statistical Analysis

Significant differences between the means plus or minus standard deviations (SDs) of different groups were examined using a two-tailed unpaired Student’s *t* test. A P value of less than 0.05 was regarded as statistically significant. All variables remained significant after multiple testing.

## ACKNOWLEDGMENTS

We thank Robert Ryan, Max Dow, Liang Yang and Delphine Caly for initial data, helpful discussions and critical reading of the manuscript. Ian Gilbert and Claire Mackenzie who provided help in resynthesizing several DSF analogues when required. This work was supported by The Wellcome Trust grant 100204/A/12/Z.

## AUTHOR CONTRIBUTIONS

SA designed the study. SA, JT, KBT and TPOS designed methods and experiments, carried out the laboratory experiments, analyzed the data, interpreted the results and wrote the paper. JT, JM, TPOS, MKG were involved in the synthesis of DSF analogues, worked on associated data collection and their interpretation. SA and KBT performed microarray and qPCR experiments and interpreted data. SA, KBT and RI designed biofilm and infection experiments, discussed analyses, interpretation, and presentation. MAV contributed to the editing and writing of the article. All authors have contributed with edits, seen and approved the manuscript.

## ACCESSION CODES

The microarray data were deposited in the Gene Expression Omnibus (GEO) database with series record GSE110126.

## REFERENCES

1. Levy SB, Marshall B (2004) Antibacterial resistance worldwide: causes, challenges and responses. Nat Med 10: S122–S129.

2. Taubes G (2008) The bacteria fight back. Science 321: 356–361.

3. Sleator RD, Hill C (2008) Battle of the bugs. Science 321: 1294–1295.

4. Sperandio V (2007) Novel approaches to bacterial a infection therapy by interfering with bacteria-to-bacteria signaling. Expert Rev Anti Infect Ther 5: 271–276.

5. Curtis MM, Russell R, Moreira CG, Adebesin AM, Wang C, Williams NS, Taussig R, Stewart D, Zimmern P, Lu B, Prasad RN, Zhu C, Rasko DA, Huntley JF, Falck JR, Sperandio V (2014) QseC inhibitors as an antivirulence approach for Gram-negative pathogens. mBIO 5.

6. Bolitho ME, Perez LJ, Koch MJ, Ng W-L, Bassler BL, Semmelhack MF. (2011) Small molecule probes of the receptor binding site in the Vibrio cholerae CAI-1 quorum sensing circuit. Bioorg. Med. Chem 19: 6906–6918.

7. O’Loughlin CT, Miller LC, Siryaporn A, Drescher K, Semmelhack MF, Bassler BL. (2013) A quorum-sensing inhibitor blocks Pseudomonas aeruginosa virulence and biofilm formation. Proc Natl Acad Sci U S A. 110(44):17981–6.

8. Williams P, Camara M (2009) Quorum sensing and environmental adaptation in Pseudomonas aeruginosa: a tale of regulatory networks and multifunctional signal molecules. Curr Opin Microbiol 12:182–191.

9. Chugani S, Greenberg EP (2014) An evolving perspective on the Pseudomonas aeruginosa orphan quorum sensing regulator QscR. Front Cell Infect Microbiol 4.

10. Jimenez PN, Koch G, Thompson JA, Xavier KB, Cool RH, Quax WJ (2012) The multiple signaling systems regulating virulence in Pseudomonas aeruginosa. Mol Biol Rev 76: 46–65.

11. Bjarnsholt T, Tolker-Nielsen T, Hoiby N, Givskov M (2010) Interference of Pseudomonas aeruginosa signalling and biofilm formation for infection control. Expert Rev Mol Med 12.

12. Jakobsen TH, van Gennip M, Phipps RK, Shanmugham MS, Christensen LD, Alhede M, Skindersoe ME, Rasmussen TB, Friedrich K, Uthe F, Jensen PO, Moser C, Nielsen KF, Eberl L, Larsen TO, Tanner D, Hoiby N, Bjarnsholt T, Givskov M (2012) Ajoene, a sulfur-rich molecule from Garlic, inhibits genes controlled by quorum sensing. Antimicrob Agents Chemother 56: 2314–2325.

13. Kim H-S, Lee S-H, Byun Y, Park H-D (2015) 6-Gingerol reduces Pseudomonas aeruginosa biofilm formation and virulence via quorum sensing inhibition. Sci Rep 5.

14. Ryan RP, Fouhy Y, Garcia BF, Watt SA, Niehaus K, Yang L, Tolker-Nielsen T, Dow JM (2008) Interspecies signalling via the Stenotrophomonas maltophilia diffusible signal factor influences biofilm formation and polymyxin tolerance in Pseudomonas aeruginosa. Mol Micro 68: 75–86.

15. Ryan RP, Dow JM (2011) Communication with a growing family: diffusible signal factor (DSF) signaling in bacteria. Trend Microbiol 19: 145–152.

16. Twomey KB, O’Connell OJ, McCarthy Y, Dow JM, O’Toole GA, Plant BJ, Ryan RP (2012) Bacterial cis-2-unsaturated fatty acids found in the cystic fibrosis airway modulate virulence and persistence of Pseudomonas aeruginosa. ISME J 6: 939–950.

17. Beaulieu ED, Ionescu M, Chatterjee S, Yokota K, Trauner D, Lindow S (2013) Characterization of a diffusible signaling factor from Xylella fastidiosa. MBIO 4.

18. Ryan RP, Dow JM (2008) Diffusible signals and interspecies communication in bacteria. Microbiol 154: 1845–1858.

19. Anderson GG, Moreau-Marquis S, Stanton BA, O’Toole GA (2008) In vitro analysis of tobramycin-treated Pseudomonas aeruginosa Biofilms on cystic fibrosis-derived airway epithelial cells. Infect Immun 76: 1423–1433.

20. Geller DE, Pitlick WH, Nardella PA, Tracewell WG, Ramsey BW (2002) Pharmacokinetics and bioavailability of aerosolized tobramycin in cystic fibrosis. Chest 122: 219–226.

21. Deng Y, Wu J, Eberl L, Zhang L-H (2010) Structural and functional characterization of diffusible signal factor family quorum-sensing signals produced by members of the Burkholderia cepacia complex. Appl Environ Microbiol. 76: 4675–4683.

22. Wang LH, He YW, Gao YF, Wu JE, Dong YH, He CZ, Wang SX, Weng LX, Xu JL, Tay L, Fang RX, Zhang LH (2004) A bacterial cell-cell communication signal with cross-kingdom structural analogues. Mol Microbiol 51: 903–912.

23. McCarthy Y, Yang L, Twomey KB, Sass A, Tolker-Nielsen T, Mahenthiralingam E, Dow JM, Ryan RP (2010) A sensor kinase recognizing the cell-cell signal BDSF (cis-2-dodecenoic acid) regulates virulence in Burkholderia cenocepacia. Mol Microbiol 77: 1220–1236.

24. Cai Z, Yuan ZH, Zhang H, Pan Y, Wu Y, Tian XQ, Wang FF, Wang L, Qian W. (2017) Fatty acid DSF binds and allosterically activates histidine kinase RpfC of phytopathogenic bacterium Xanthomonas campestris pv. campestris to regulate quorum-sensing and virulence. PLoS Pathog. 2017 Apr 3;13(4):e1006304. doi: 10.1371/journal.ppat.1006304.

25. Ng W-L, Wei Y, Perez LJ, Cong J, Long T, Koch M, Semmelhack MF, Wingreen NS, Bassler BL (2010) Probing bacterial transmembrane histidine kinase receptor-ligand interactions with natural and synthetic molecules. Proc Natl Acad Sci USA 107:5575–5580.

26. Deng Y, Lim A, Lee J, Chen S, An S, Dong Y-H, Zhang L-H (2014) Diffusible signal factor (DSF) quorum sensing signal and structurally related molecules enhance the antimicrobial efficacy of antibiotics against some bacterial pathogens. BMC Microbiol 14.

27. Vonheijne G (1992) Membrane-protein structure prediction - Hydrophobicity analysis and the positive-inside rule. J Mol Biol 225: 487–494.

28. Sonnhammer EL, von Heijne G, Krogh A (1998) A hidden Markov model for predicting transmembrane helices in protein sequences. Intel Sys Mol Biol 6:175–182.

29. Cserzo M, Eisenhaber F, Eisenhaber B, Simon I (2002) On filtering false positive transmembrane protein predictions. Protein Eng 15: 745–752.

30. Cserzo M, Wallin E, Simon I, vonHeijne G, Elofsson A (1997) Prediction of transmembrane alpha-helices in prokaryotic membrane proteins: the dense alignment surface method. Protein Eng 10: 673–676.

31. Hirokawa T, Boon-Chieng S, Mitaku S. (1998) SOSUI: classification and secondary structure prediction system for membrane proteins. Bioinfo 14:378–379.

32. Tusnady GE, Simon I (2001) The HMMTOP transmembrane topology prediction server. Bioinfo 17: 849–850.

33. Daniels C, Vindurampulle C & Morona R (1998) Overexpression and topology of the Shigella flexneri O-antigen polymerase (Rfc/Wzy). Mol Microbiol 28: 1211–1222.

34. Baker SJ, Daniels C & Morona R (1997) PhoP/Q regulated genes in Salmonella typhi identification of melittin sensitive mutants.Microb Pathogen 22: 165–179

35. Tidwell TT (1990) Oxidation of alcohols by activated dimethyl-sulfoxide and related reactions - an update. Synthesis 5:857–870.

36. Takeda T (2004) Modern Carbonyl Olefination: Methods and Applications. Wiley.

37. Dayal B, Salen G, Tint GS, Shefer S, Benz SW (1990) Use of positive ion fast atom bombardment mass spectrometry for rapid identification of a bile alcohol glucuronide isolated from cerebrotendinous Xanthomatosis patients. Steroids 55:74–78.

38. Sivasubramanian K, Kaanumalle LS, Uppili S, Ramamurthy V (2007) Value of zeolites in asymmetric induction during photocyclization of pyridones, cyclohexadienones and naphthalenones. Org Biomol Chem 5: 1569–1576.

39. Montalbetti C, Falque V (2005) Amide bond formation and peptide coupling. Tetrahedron 61: 10827–10852.

40. Barber CE, Tang JL, Feng JX, Pan MQ, Wilson TJG, Slater H, Dow JM, Williams P, Daniels MJ (1997) A novel regulatory system required for pathogenicity of Xanthomonas campestris is mediated by a small diffusible signal molecule. Mol Microbiol 24: 555–566.

41. Clarke MB, Hughes DT, Zhu C, Boedeker EC, Sperandio V (2006) The QseC sensor kinase: A bacterial adrenergic receptor. Proc Natl Acad Sci USA 103:10420–10425.

42. Rigaud JL, Levy D (2003) Reconstitution of membrane proteins into liposomes. Liposomes 372:65–86.

43. McCaughey LC, Ritchie ND, Douce GR, Evans TJ, Walker D (2016) Efficacy of species-specific protein antibiotics in a murine model of acute Pseudomonas aeruginosa lung infection. Sci Rep. 6:30201.

44. An S-Q, Caly DL, McCarthy Y, Murdoch SL, Ward J, Febrer M, Dow JM, Ryan RP (2014) Novel cyclic di-GMP effectors of the YajQ protein family control bacterial virulence. Plos Pathogens 10: e1004429.

45. An S-Q, Febrer M, McCarthy Y, Tang D-J, Clissold L, Kaithakottil G, Swarbreck D, Tang J-L, Rogers J, Dow JM, Ryan RP (2013) High-resolution transcriptional analysis of the regulatory influence of cell-to-cell signalling reveals novel genes that contribute to Xanthomonas phytopathogenesis. Mol Micro 88:1058–1069.

